# Risky behaviors and Parkinson’s disease: A Mendelian randomization study in up to 1 million study participants

**DOI:** 10.1101/446807

**Authors:** Sandeep Grover, Greco M Fabiola Del, Meike Kasten, Christine Klein, Christina M. Lill, Inke R. König

**Affiliations:** Institut für Medizinische Biometrie und Statistik, Universität zu Lübeck, Universitätsklinikum Schleswig-Holstein, Campus Lübeck, Lübeck, Germany; Institute for Biomedicine, Eurac research, Via Galvani 31, Bolzano, Italy; Institute of Neurogenetics, Department of Psychiatry and Psychotherapy, Universität zu Lübeck, Lübeck, Germany; Institute of Neurogenetics, Universität zu Lübeck, Lübeck, Germany; Genetic and Molecular Epidemiology Group, Lübeck Interdisciplinary Platform for Genome Analytics (LIGA), Institutes of Neurogenetics & Cardiogenetics, Universität zu Lübeck, Lübeck, Germany

## Abstract

**Objective:** Dopaminergic neurotransmission is known to be a potential modulator of risky behaviors including substance abuse, promiscuity, and gambling. Furthermore, observational studies have shown associations between risky behaviors and Parkinson’s disease; however, the causal nature of these associations remains unclear. Thus, in this study, we examine causal associations between risky behavior phenotypes on Parkinson’s disease using a Mendelian randomization approach.

**Methods:** We used two-sample Mendelian randomization to generate unconfounded estimates using summary statistics from two independent, large meta-analyses of genome-wide association studies on risk taking behaviors (n=370,771-939,908) and Parkinson’s disease (cases: n=9581, controls: n = 33,245). We used inverse variance weighted as the main method for judging causality.

**Results:** Our results support a strong protective association between the tendency to smoke and Parkinson’s disease (OR=0.714 per log odds of ever smoking; 95% CI=0.568-0.897; p-value=0.0041; Cochran Q test; p-value=0.238; I^2^ index=6.3%). Furthermore, we observed risk association trends between automobile speed propensity as well as the number of sexual partners and Parkinson’s disease after removal of overlapping loci with other risky traits (OR=1.986 for each standard deviation increase in normalized automobile speed propensity; 95% CI=1.215-3.243; p-value=0.0066, OR=1.635 for each standard deviation increase in number of sexual partners; 95% CI=1.165-2.293; p-value=0.0049).

**Interpretation:** These findings provide support for a causal relationship between general risk tolerance and Parkinson’s disease and may provide new insights in the pathogenic mechanisms leading to the development of Parkinson’s disease.

## Introduction

Parkinson’s disease (PD) is the second most prevalent neurodegenerative disorder characterized pathologically by progressive loss of dopaminergic neurons in the substantia nigra ^1^. The currently available treatment options for PD are symptomatic only. The lack of disease-modifying or protective treatments is at least in part due to the fact that the exact disease mechanisms are currently only partly understood.

The vast majority of PD cases are caused by the combined action and likely interaction of genetic variants as well as environmental and lifestyle exposures ^2–6^. Several common habitual agents like smoking, coffee and alcohol drinking have shown protective associations with PD in large scale meta-analyses of observational studies ^7^. Several recent studies have further shown beneficial effects of cannabidiol, a non-psychotomimetic compound derived from cannabis on non-motor symptoms in PD patients ^8^. It is noteworthy that several impulse control disorders (ICDs) such as gambling, hypersexuality and compulsive eating are observed more frequently in PD patients with some studies reporting up to 40% prevalence of ICDs in PD patients^9^. However, it is believed that these symptoms may be the result of dopamine agonist therapy prescribed to PD patients^10^. The imminent challenge in this context is to decipher whether these PD-associated environmental/lifestyle/behavioral variables contribute to or are an effect of the disease.

The recent development of the Mendelian randomization (MR) approach allows to judge causality based on genetic data generated in observational studies. Specifically, this relies on the utilization of genetic variants as proxy markers of risk factors ^11^ and takes care of confounding by exploiting random allocation of genetic variants at birth. We have also seen a surge of MR studies in the field of PD specifically exploring the causal role of several circulating biomarkers ^12–18^. For example, a recent MR study reported a significant causal association with a lifelong PD risk reduction of 3% per 10 μg/dl increase in serum iron levels ^14^. Most recently, another study further reported a risk reduction of 18% with a lifetime exposure of 5kg/m2 higher BMI ^15^.

To date, the majority of studies in the field of PD have focused on modifiable environmental factors only, and MR studies exploring the role of behavioral phenotypes are lacking. Henceforth, our primary aim was to investigate the willingness to take risk as a causal factor in the development of PD. For this, we applied a two-sample MR to investigate whether people with risk taking tendency have an altered risk for PD ^19^. A recent GWAS identified 611 independent loci with several measures of risky behaviors including general risk tolerance, adventurousness, and risky behaviors in the driving, drinking, smoking, and sexual domains ^20^. We used all reported loci to mimic the random allocation of loci among PD cases and controls in available data from a large, recent GWAS on PD ^4, 21^. As secondary analyses, we further considered the wider literature to support inferences drawn from our primary analysis using previously reported GWAS on similar habitual behaviors such as smoking phenotypes, coffee consumption, alcoholism, cannabis dependence and gambling ^22–27^.

## Methods

### Study design and identification of datasets

We conducted a two-sample MR using summary level estimates to explore the causal role of several risky behaviors on PD ^28^. We identified genetic instruments that influence risky behaviors using a recently published meta-analysis of GWAS datasets on risky behaviors ^20^. The study reported statistically significant associations of 611 independent loci (p-value<5×10^−8^) in a discovery cohort in up to 939,908 individuals of European ancestry with six highly correlated risky behavior phenotypes including general risk tolerance, adventurousness, automobile speeding propensity, drinks per week, ever versus never smoking and number of sexual partners. The study further defined general risk tolerance as the willingness to take risks, “adventurousness” as the self reported tendency to be adventurous vs. cautious, “automobile speeding propensity” as the tendency to drive faster than the speed limit, “drinks per week” as the average number of alcoholic drinks consumed per week, “ever smoker (tendency to smoke)” as whether one has ever been a smoker, and lastly “number of sexual partners” as the lifetime number of sexual partners.

We further extracted summary estimates of the identified genetic variants from the discovery cohort of a recent meta-analysis of GWAS on 9581 PD cases and 33, 245 controls of European ancestry ^4^. For this, we used data available on the PDGene database (http://www.pdgene.org)^21^. Genetic instruments were identified for smoking (cigarettes per day), smoking initiation, smoking cessation, cannabis dependence, pathological gambling, alcohol and coffee consumption from independent GWAS as a part of our secondary analyses ^22–27^.

### Prioritization of genetic variants and power analysis

We systematically screened all the identified loci for a possible direct involvement in PD. For this, we used data available via PDGene to extract the list of loci shown to be significantly associated with PD (p-value<5×10^−8^) ^21^. We further checked overlapping loci for a relevant role in the pathogenesis of PD using a literature search. If substantial evidence was found, the respective loci were excluded from the list of genetic variants (SNPs, genetic instrument) of the respective behavioral traits.

The SNPs constituting each genetic instrument were checked for strong linkage disequilibrium (LD). We used the rAggr database to look for correlated variants in individuals with European descent from the 1000 Genomes Phase 3 data (http://raggr.usc.edu; date last accessed June 22, 2018) and excluded one of the variants for pairs with R^2^ greater than 0.25. Finally, if SNPs were not available in the PD GWAS dataset, we identified proxy SNPs using an R^2^ cut-off of 0.9 based on the rAggr database as above.

The strength of the prioritized genetic instrument was judged using F-statistics as explained earlier^14^. We computed the variance in exposures explained by prioritized genetic instruments (R^2^) of the genetic insturments using effect estimates and the standard error of individual SNPs as described elsewhere^29^. Lastly, power calculations were done using the method described by Brion *et al.*, which is available online http://cnsgenomics.com/shiny/mRnd/) ^30^.

### Estimation of causal effects

In cases where the genetic instruments comprised a single SNP, we used the Wald ratio estimate along with the Delta method to obtain the related estimate of the variance. In cases where the genetic instruments consisted of multiple SNPs, we used the inverse variance weighted (IVW) fixed effect method as the main method to estimate the effect of genetically predicted behavioral phenotypes on PD by combining the genetic loci-specific Wald ratio estimates. We specifically employed the IVW method using second order weights because casual estimates generated through this method are expected to provide a more accurate reflection of the variance of the Wald ratio estimate ^31^.

However, in the absence of reliable information on functional pathways, proportion and direction of pleiotropic genetic variants, additional MR methods including MR-Egger, Weighted median and Weighted mode-based method were also employed to check the consistency of direction of effect estimates ^19, 32–34^. Unlike IVW, which assumes no intercept term in the model, the MR-Egger method provides less biased causal estimates in the presence of directional pleiotropy and considerable heterogeneity assuming absence of measurement error (NOME assumption) ^19^. However, the MR-Egger method is more sensitive to unobserved associations of genetic variants with confounders of the exposure-outcome association and requires a greater sample size for the same underlying variance in exposure ^33^. Both IVW and MR-Egger methods further assume that the pleiotropic effects of genetic variants are independent of their associations with the exposure known as the InSIDE assumption. In the case of violation, the Weighted median method may provide consistent causal estimates even if up to 50% of genetic variants do not conform to the InSIDE assumption. Also the Weighted mode-based method may provide consistent causal estimates, in particular, even when the NOME assumption was not met, but assuming that the most frequent value of the bias of the Wald ratio estimates is zero. ^34^

Within every MR method, we computed casual estimates as odds ratio (OR) for PD per unit log of odds of the categorical behavioral phenotypes or OR per unit standard deviation (SD) of the continuous behavioral phenotypes. And lastly, to address the issue of multiple testing, results were considered statistically significant at the 5% level after a conservative Bonferroni correction of the significance level, therefore if p-value<8.3×10^−3^ (0.05/6 independent primary MR hypotheses).

### Assessment of pleiotropy

We used the Cochran Q-statistic and I^2^ for the IVW method using second order weights as main methods to identify pleiotropic variants ^35^. Furthermore, results from the less powerful MR-Egger’s test were also used to explore heterogeneity including the test for deviation of the intercept from the null for MR-Egger’s model using the χ^2^-test for independence ^33^. We further used Ruckers Q’ statistic to describe heterogeneity around MR-Egger fit ^36^. The appropriate use of the main MR method for interpretation of causal estimate in the present study was judged by calculating the ratio between Rucker’s Q’ and Cochran’s Q statistics ^37^. As a rule of thumb, the IVW method is recommended as the main method for judging the causal effect if the ratio approaches one.

To evaluate heterogeneity graphically, funnel plots were constructed that plot the spread of the inverse of the standard error of the respective Wald ratio estimates of each individual SNP around the MR estimates. Also, scatter plots of effect estimates of individual SNPs with outcome vs. effect estimates of individual SNPs with exposure are provided as a comparative visual assessment of the causal estimates generated from different MR methods. We further constructed radial MR plots which have been recently suggested as a more suitable approach for visual detection of outliers compared to traditional scatter plots, specifically when the difference between IVW and MR-Egger estimates is large ^38^.

### Sensitivity analyses

A leave-one-out sensitivity analysis was conducted to check for a disproporationate influence of individual SNPs on overall causal effect estimates using the IVW method. We used forest plots to visually assess the results of the analysis and further identify the outliers.

Since all the behavioral traits are highly correlated and are expected to exhibit shared genetic influence, we conducted a sensitivity analysis by including only genetic loci specific to each individual behavioral trait. We used an R^2^>0.8 to consider loci to be overlapping with other loci in an independent genetic instrument. Such an approach may help us to judge the reliability of independent associations of observed phenotypic traits. We further adopted a conservative approach by using loci unique to each phenotypic trait (R^2^<0.01) at the cost of reduced power.

We used Phenoscanner database to identify potential pleiotropic variants by checking significant associations of loci prioritized in the present study with phenotypes from previously published GWAS (http://phenoscanner.medschl.cam.ac.uk) ^39^. We further checked GWAS listed in the GWAS Catalog (https://www.ebi.ac.uk/gwas/) to search for any missed hits. The identified variants were then grouped into categories depending on their association with potential confounder phenotypes and were checked for an influence on the causal estimate using leave-out approach.

We further evaluated the biological relevance of different brain regions in contribution to the overall causal estimate through analysis of gene expression data for the available loci from our different genetic instruments. Gene expression data was extracted from the Genotype-Tissue Expression (GTEx) Project comprising data on a total of 12 different brain regions (www.gtexportal.org; date last accessed June 22, 2018) ^40^. The identified loci were then grouped into categories as per their expression in specific brain regions and checked for an influence on the causal estimate after their exclusion using a leave-out approach.

## Results

### Prioritization of genetic instruments and power analysis

The descriptive statistics of the genetic instruments selected for the MR analyses are presented in **Table 1**. The data used for the analyses are given in the **Supplementary Table**.

**Table 1.** Summary of genetic instruments used in the present Mendelian randomization analysis. PD = Parkinson’s disease Maximum sample size in PD cohort = 42,286 individuals. *p-value<5×10^−8^ for association with risky behavior in GWAS (20) **Excluded from further analysis (identified using pd.org database) Automobile speed propensity: MAPT (rs62062288); HLA-DQB (rs3021058) Drinks per week: MAPT (rs62055546) Number of sexual partners: MAPT (rs62063281) ***Excluded from further analysis Adventurousness: rs1492436 and rs35377646 (r2=1.0 with similar effect estimates on exposure). rs1492436 was selected for the present study based on better variance explained by the SNP

| Risky behavior | Maximum sample size (N) | # SNPs associated with risky behavior* | # SNPs with direct influence on PD ** | # SNPs in high LD ($R^2>0.25$)*** | # proxy SNPs ($R^2>0.9$) | # SNPs in genetic instrument | F-statistic |
| --- | --- | --- | --- | --- | --- | --- | --- |
| General risk tolerance | 939,908 | 124 | 0 | 0 | 10 | 117 | 5064.4 |
| Adventurousness | 557,923 | 167 | 0 | 1 | 16 | 150 | 6874.5 |
| Automobile speeding propensity | 404,291 | 42 | 2 | 0 | 2 | 35 | 1406.4 |
| Drinks per week | 414,343 | 85 | 1 | 0 | 9 | 70 | 4342.4 |
| Tendency to smoke | 518,633 | 223 | 0 | 0 | 11 | 213 | 9639.6 |
| Number of sexual partners | 370,711 | 118 | 1 | 0 | 12 | 109 | 4515.6 |

Two SNPs from the genes *MAPT* (rs62062288; p-value with PD=3.1 x 10^−21^) and *HLA-DQB* (rs3021058; p-value with PD= 2.7 x 10^−3^) were excluded from the genetic instrument for automobile speeding propensity phenotype based on a potential direct involvement in PD. Two additional *MAPT* SNPs were also present in the respective genetic instrument for the phenotypes drinks per week (rs62055546; p-value with PD: 5.2 x 10^−21^) and number of sexual partners (rs62063281; p-value with PD=1.73 2 x 10^−21^) and were not carried forward to further analyses. One SNP was observed to be in complete LD with another SNP for the adventurousness phenotype and was excluded. The final number of available SNP data varied from 35 (out of 42) for automobile speeding propensity to 213 (out of 223) for smoking tendency with F-statistics of the pooled genetic instrument ranging from 1406.4 (for automobile speeding propensity) to 9639.6 (for ever vs. never smoking).

Our power analyses suggest that our study has approximately 80% power to detect a true OR of 1.349 or 0.698 for PD per SD of the continuous phenotype assuming that the proportion of the continuous phenotype explained by the genetic instrument is ≥1% at a type-1 error rate of 0.05.

### Estimation of causal effects and assessment of pleiotropy

The causal effect estimates using different MR methods are provided in **Table 2**. Using the IVW method, a genetically increased risk of tendency to smoke was associated with a reduced risk of PD per unit increase in log odds of ever smoking (OR: 0.714 per log odds of ever smoking; 95% CI=0.568-0.897; p-value=0.0041). Results from the Weighted median MR analysis showed similar results (OR: 0.707 per log odds of ever smoking; 95% CI=0.601-0.832). There was minimal evidence of heterogeneity of causal effects between individual variants (I^2^ = 6.30%; Cochran’s Q test p-value=0.2367), which was confirmed using MR-Egger’s Intercept test (p-value=0.6619). Corresponding plots used for the assessment of pleitropy are shown in **Figure 1**. We did not detect any potential outlier or pleiotropic variant in the association analysis of the tendency to smoke phenotype and PD.

**Table 2.** Causal effect estimates using different Mendelian randomization methods and heterogeneity analysis of causal effect estimates for risk taking behaviors. P-values for effects on PD marked in bold show statistical significance after Bonferroni corrections with a cut-off p-value of 0.05/6 = 0.0083 P-values for test on heterogeneity marked in bold are below 0.05. General risk tolerance: OR per log odds of general risk tolerance Adventuorness: OR per log odds of adventourness Automobile speeding propensity: OR per SD of normalized Automobile speeding propensity Drink per weeks: OR per SD of number of drinks per week Ever smokers: OR per log odds of ever smoking Number of sexual partners: OR per SD of number of sexual partners PD = Parkinson’s disease * Computed using second order weights

| Risky behavior | MR methodology | Causal effect estimates<br>(PD as outcome) |  |  | Tests of heterogeneity |  |
| --- | --- | --- | --- | --- | --- | --- |
|  |  | OR* | 95%CI | P-value | Test | P-value |
| General risk tolerance | Inverse variance weighted (2nd order weights) | 1.620 | 1.046-2.511 | 0.0311 | MR-Egger intercept (p-value) | 0.2114 |
|  | MR-Egger | 0.406 | 0.042-3.934 | 0.4335 | I<sup>2</sup> (%) | 8.34% |
|  | Weighted median method | 1.122 | 0.820-1.535 | 0.7148 | Cochrane Q-test (p-value) | 0.2367 |
| | Weighted mode method (NOME assumptions) | 0.708 | 0.141-3.550 | 0.6751 | Rucker's Q$\chi^2$-test (p-value) | 0.2507 |
| | | | | | Rucker's Q$\chi^2$ statistic /Cochrane Q statistic | 1.0141 |
| Adventurousness | Inverse variance weighted (2nd order weights) | 1.091 | 0.810-1.470 | 0.5620 | MR-Egger intercept (p-value) | 0.0796 |
|  | MR-Egger | 0.382 | 0.111-1.318 | 0.1192 | I<sup>2</sup> (%) | 15.85% |
|  | Weighted median method | 0.879 | 0.712-1.085 | 0.5427 | Cochrane Q-test (p-value) | 0.0579 |
| | Weighted mode method (NOME assumptions) | 0.607 | 0.252-1.460 | 0.2667 | Rucker's Q$\chi^2$-test (p-value) | 0.0730 |
| | | | | | Rucker's Q$\chi^2$ statistic /Cochrane Q statistic | 1.0194 |
| Automobile speeding propensity | Inverse variance weighted (2nd order weights) | 2.043 | 1.076-3.876 | 0.0299 | MR-Egger intercept (p-value) | 0.0617 |
|  | MR-Egger | 0.125 | 0.006-2.654 | 0.1757 | I<sup>2</sup> (%) | 28.75% |
|  | Weighted median method | 2.738 | 1.856-4.040 | 0.0140 | Cochrane Q-test (p-value) | 0.0594 |
| | Weighted mode method (NOME assumptions) | 5.202 | 0.673-40.193 | 0.1232 | Rucker's Q$\chi^2$-test (p-value) | 0.1213 |
| | | | | | Rucker's Q$\chi^2$ statistic /Cochrane Q statistic | 1.1189 |
| Drinks per week | Inverse variance weighted (2nd order weights) | 1.150 | 0.868-1.525 | 0.3247 | MR-Egger intercept (p-value) | **0.0408** |
|  | MR-Egger | 0.791 | 0.498-1.257 | 0.3159 | I<sup>2</sup> (%) | 0.00% |
|  | Weighted median method | 0.845 | 0.676-1.055 | 0.4513 | Cochrane Q-test (p-value) | 0.6032 |
| | Weighted mode method (NOME assumptions) | 0.855 | 0.543-1.344 | 0.4992 | Rucker's Q$\chi^2$-test (p-value) | 0.7071 |
| | | | | | Rucker's Q$\chi^2$ statistic /Cochrane Q statistic | 1.0672 |
| Tendency to smoke | Inverse variance weighted (2nd order weights) | 0.714 | 0.568-0.897 | **0.0041** | MR-Egger intercept (p-value) | 0.6619 |
|  | MR-Egger | 0.552 | 0.185-1.646 | 0.2849 | I<sup>2</sup> (%) | 6.30% |
|  | Weighted median method | 0.707 | 0.601-0.832 | 0.0339 | Cochrane Q-test (p-value) | 0.2389 |
| | Weighted mode method (NOME assumptions) | 0.242 | 0.083-0.703 | **0.0098** | Rucker's Q$\chi^2$-test (p-value) | 0.2276 |
| | | | | | Rucker's Q$\chi^2$ statistic /Cochrane Q statistic | 1.0010 |
| Number of sexual partners | Inverse variance weighted (2nd order weights) | 1.473 | 1.079-2.010 | 0.0152 | MR-Egger intercept (p-value) | 0.3621 |
|  | MR-Egger | 0.674 | 0.113-4.006 | 0.6615 | I<sup>2</sup> (%) | 21.22% |
|  | Weighted median method | 1.365 | 1.103-1.688 | 0.1465 | Cochrane Q-test (p-value) | **0.0308** |
| | Weighted mode method (NOME assumptions) | 1.270 | 0.446-3.618 | 0.6547 | Rucker's Q$\chi^2$-test (p-value) | 0.0298 |
| | | | | | Rucker's Q$\chi^2$ statistic /Cochrane Q statistic | 1.0064 |

**Figure 1:**
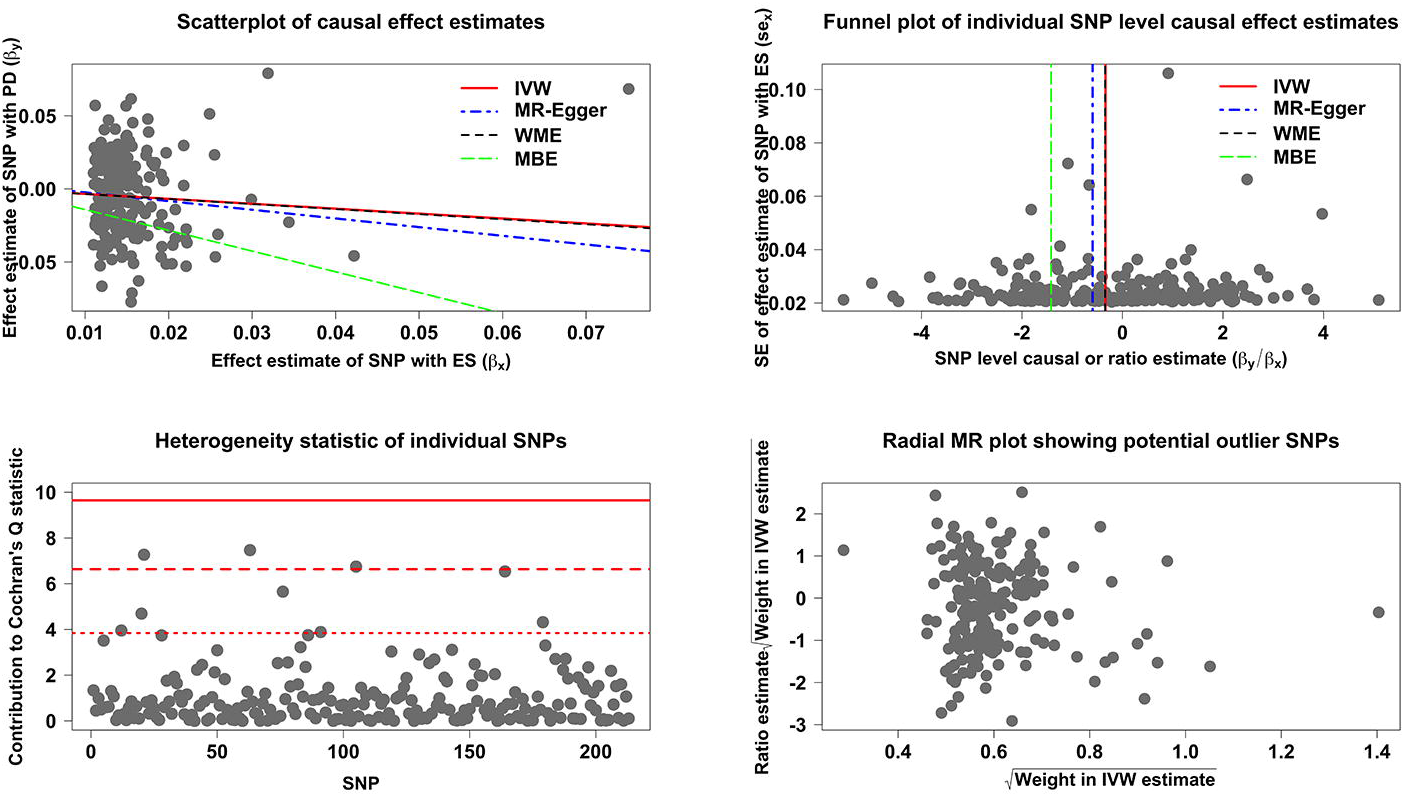
Causal association analysis and assessment of pleiotropy for the ever smoking phenotype with Parkinson’s disease A. Scatterplot showing causal effect estimates computed using various MR methods. B. Funnel plot showing the extent of heterogeneity among the individual Wald ratio estimates. C. Plot of Cochran’s Q estimates for individual SNPs constituting the genetic instrument for ever smoker phenotype using IVW method employing second order weights. D. Radial MR plot showing the distribution of weights contributed by individual SNPs in the causal effect estimation by IVW method employing second order weights.

For other risk taking behaviors including general risk tolerance, automobile speeding propensity and the number of sexual partners, we observed a trend towards positive associations (general risk tolerance: OR: 1.620 per log odds of general risk tolerance; 95% CI=1.046-2.511; p-value=0.0311; automobile speeding propensity: OR=2.043 for each SD increase in normalized automobile speed propensity; 95%CI=1.076-3.876; p-value=0.0299; number of sexual partners: OR: 1.473 for each SD increase in the number of sexual partners; 95%CI=1.079-2.010; p-value=0.0152). A triangulation flowchart summarizing the findings of the study in the context of the MR workflow is given in **Figure 2**.

**Figure 2:**
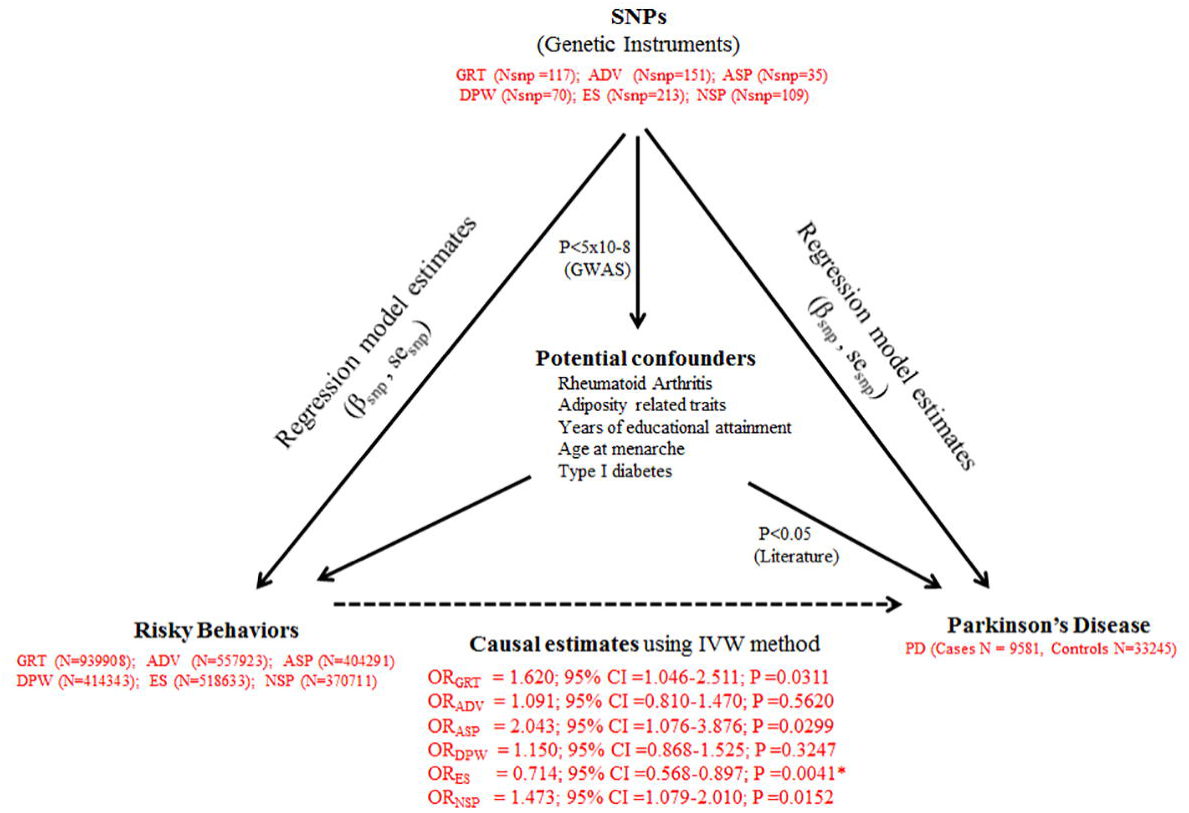
Triangular representation of results from the present MR study Abbreviations in the figure: SNP: Single Nucleotide Polymorphism Nsnp: Number of SNPs in the genetic instrument for each respective risky behavior β_snp_: Regression coeffiecient corresponding to each specific SNP for respective arm of the triangle corresponding to each relationship se_snp_: Standard error estimates corresponding to each specific SNP for the respective arm of the triangle corresponding to each relationship GRT: General risk tolerance ADV: Adventurousness ASP: Automobile speeding propensity DPW: Drinks per week ES: Ever vs never smoking NSP: Number of sexual partners OR_GRT_, OR_ADV_, OR_ES_: Expressed as odds ratio per log odds of respective risky behavior OR_ASP_, OR_DPW_, OR_NSP_: Expressed as odds ratio per SD of respective risky behavior

### Sensitivity analyses

#### Leave-one-out analysis

Based on the forest plots, we observed outlier SNPs for the phenotypes general risk tolerance (rs993137), adventurousness (rs10433500) and drinks per week (rs1229984) (data not shown). Leaving out outlier SNPs for each of the respective phenotypes did not alter the results substantially (general risk tolerance OR=1.763 per log odds of general risk tolerance; 95% CI=1.133-2.744; p-value=0.0125, adventurousness OR=1.164 per log odds of adventourness; 95% CI=0.858-1.580; p-value=0.3270, drinks per week OR=1.368 for each SD increase in the number of drinks per week; 95% CI=0.978-1.915; p-value=0.0669).

#### Genetic overlap between risky behaviors

We identified a reduction in the number of unique SNPs in the genetic instruments for each of the phenotypes using two different LD cut-offs (R^2^≤0.8 and R^2^ ≤0.01) (**Table 3**). However, irrespective of cut-off crirteria, there was no change in the protective association of the tendency to smoke phenotype with PD (number of SNPs in the genetic instrument with R^2^≤0.8: 195, OR=0.713 per log odds of ever smoking; 95%CI=0.557-0.913; p-value=0.0037; number of SNPs in the genetic instrument with R^2^ ≤0.01: OR=0.719 per log odds of ever smoking; 95%CI=0.547-0.945; p-value=0.0185). Furthermore, consistent with the risk-increasing trend observed for general risk tolerance and the number of sexual partners, we observed a stronger causal association with PD for both phenotypes after reducing the number of SNPs from 117 to 94 (R^2^≤0.8) for general risk tolerance and 109 to 94 (R^2^≤0.8) for the number of sexual partners (OR=1.986 per log odds of general risk tolerance; 95% CI=1.215-3.243; p-value= 0.0066, OR=1.635 for each SD increase in number of sexual partners; 95% CI=1.165-2.293; p-value=0.0049). The associations persisted using a stringent lower R^2^ cut-off of 0.01 (p-value=0.0440 and p-value=0.0484).

**Table 3.** Causal effect estimates using unique loci among different phenotypic traits. PD = Parkinson’s disease * Computed using second order weights

| Risky behavior | MR methodology | Number of SNPs ($R^2 \leq 0.80$ across phenotypes) | Causal effect estimates (PD as outcome) | | | Number of SNPs ($R^2 \leq 0.01$ across phenotypes) | Causal effect estimates (PD as outcome) | | |
| --- | --- | --- | --- | --- | --- | --- | --- | --- | --- |
|  |  |  | OR (95% )* | 95%CI | p-value |  | OR (95% )* | 95%CI | p-value |
| General risk tolerance | Inverse variance weighted* | 94 | 1.986 | 1.215-3.243 | **0.0066** | 66 | 1.821 | 1.017-3.261 | **0.0440** |
| Adventurousness | Inverse variance weighted* | 125 | 1.169 | 0.837-1.633 | 0.3564 | 80 | 1.002 | 0.672-1.493 | 0.9915 |
| Automobile speeding propensity | Inverse variance weighted* | 31 | 2.279 | 1.130-4.597 | **0.0229** | 19 | 2.498 | 0.893-6.992 | 0.0780 |
| Drinks per week | Inverse variance weighted* | 67 | 1.094 | 0.819-1.459 | 0.5370 | 52 | 1.091 | 0.800-1.488 | 0.5755 |
| Ever smoker | Inverse variance weighted* | 195 | 0.713 | 0.557-0.913 | **0.0075** | 154 | 0.719 | 0.547-0.945 | **0.0185** |
| Number of sex partners | Inverse variance weighted* | 94 | 1.635 | 1.165-2.293 | **0.0049** | 50 | 1.585 | 1.003-2.502 | **0.0484** |

#### Geneticvariants associated with potential confounders

We did a comprehensive screening of the the Phenoscanner database for potential associations of genetic loci used in the current study and reported to be associated with other phenotypes. The identified associated phenotypes were then investigated for association with PD based on a thorough literature search. Using this strategy, we identified rheumatoid arthritis, years of educational attainment, adiposity related traits, age at menarche and type I diabetes as potential confounders ^41–45^ (**Figure 2**). We identified eight genetic variants or loci from our genetic instrument for the tendency to smoke phenotype associated with different confounding traits. SNP rs12042017 has been previously reported to be associated with years of educational attainment (p-value=4.48×10^−10^). SNPs rs13396935 and rs6265 were observed to be associated with several adiposity related measures. The proxy variant rs1514174 of rs4650277 (R^2^=0.99) was further associated with BMI (p-value=2.99×10^−27^). Rheumatoid arthritis, age at menarche and type I diabetes were further identified as potential confounders associated with rs2734971, rs4650277 and rs1701704 (proxy for rs772921 with complete LD). Our sensitivity analysis excluding each of the SNPs or combinations of SNPs based on their common associated trait showed no overall influence on causal effect estimate for the tendency to smoke phenotype (data not shown).

#### Geneticvariants involved in brain expression

Using brain-specific expression quantitative trait loci (eQTL) retrieved from GTEx, we identified 27 different SNPs from the genetic instrument for the tendency to smoke phenotype with varied influence in different brain regions (data not shown). Suprisingly, the corresponding candidate genes were least represented in the substantia nigra, while as many as 10 genetic variants were observed to significantly influence gene expression in cerebellar hemisphere as well as cerebellum. Our sensitivity analysis showed that excluding genetic variants mapping to genes over-expressed in the cerebellum had maximum influence on the overall causal effect estimation (OR=0.761; 95% CI=0.606-0.957; p-value=0.0197). A similar influence was observed after excluding all the genetic variants mapping to genes expressed in brain (OR=0.735; 95% CI=0.581-0.930; p-value=0.0106). Our sensitivity analysis thereby suggested an important role of the cerebellum in the smoking tendency phenotype. Our literature search for the excluded genetic variants in the tendency to smoke genetic instrument influencing expression in cerebellum for potential influence on other biological pathways rules out pleiotropic effect of these variants.

### Secondary MR analysis

The descriptive statistics of the genetic instruments selected for the secondary MR analyses are presented in **Table 4**. The causal effect estimates are shown in **Table 5**. We first employed genetic instruments for different traits representative of the smoking phenotype (ever smoker vs. never been a regular smoker, former vs. current smoker and cigarettes per day) ^22–23^. A previous meta-analysis of GWAS on the ever smoker phenotype in 143,023 individuals of European ancestry identified genetic variants from the *BDNF* gene to be associated with the ever smoker phenotype ^22^. Since all the variants were in high LD with each other, we used rs6265, a non-synonymous variant with a functional effect on gene expression, as a proxy for all other variants for the MR analysis. We failed to observe any association (OR=0.545; 95% CI=0.230-1.291; p-value=0.1681). Interestingly, a genetic instrument (rs3025343) for former smokers vs. current smokers showed a trend towards risk for predisposition to PD (OR=1.874; 95% CI=1.003-3.499; p-value=0.0487). We further extracted a genetic instrument comprised of four uncorrelated genetic variants for a related phenotype of cigarettes per day from a meta-analysis of GWAS on 86,956 individuals, again failing to observe any trend (OR=0.989; 95% CI = 0.870-1.124; p-value=0.7995) ^23^.

**Table 4.** Summary of genetic instruments used in the Mendelian randomization analysis based on risky-and habit-related behaviors from previous GWAS in European populations. PD = Parkinson’s disease * For 1 SNP (rs4105144), no proxy was available in PD dataset **For 2 SNPs, no proxy was available *** Two different sets of SNPs were used. rs2472297 and rs2470893 showed moderate LD (r2=0.658) and hence 2 separate MR analyses were conducting including one SNP at a time.

| Habitual phenotype | Phenotypic definition | Reference | Maximum sample size | GWAS study cohort used for extraction of effect estimates | GWAS cut-off used for prioritization of SNPs (p-value) | # significant SNP/s | # proxy SNPs (R<sup>2</sup>>0.9) | # SNPs in high LD (R<sup>2</sup>>0.25) excluded | # SNPs in genetic instrument |
| --- | --- | --- | --- | --- | --- | --- | --- | --- | --- |
| Smoking | Cigarettes per day | Thorgeirsson *et al.* 2010 [23] | 86,956 | Pooled | 5x10<sup>-8</sup> | 6 | 0 | 1 | 4* |
| Smoking initiation | Ever smoker vs. never been a regular smoker | Tobacco Genetic Consortium 2010 [22] | 143,023 | Pooled | 5x10<sup>-8</sup> | 8 | 0 | 7 | 1 |
| Smoking cessation | Former vs. current smoker | Tobacco Genetic Consortium 2010 [22] | 64,924 | Pooled | 5x10<sup>-8</sup> | 1 | 0 | NA | 1 |
| Cannabis dependence | Cannabis dependence (DSMIV) vs. Individuals not meeting cannabis dependence criteria with a history of at least once in lifetime use of cannabis | Agrawal *et al.* 2018 [25] | 8515 | Discovery | 1x10<sup>-6</sup> | 26 | 0 | 25 | 1 |
| Pathological Gambling | Individuals diagnosed with pathological gambling (DSMIII/IV) vs. population controls | Lang *et al.* 2017 [26] | 1431 | Discovery | 1x10<sup>-4</sup> | 57 | 1 | 12 | 45 |
| Alcohol consumption | Units of alcohol consumed in the previous week | Clarke *et al.* 2017 [24] | 112,117 | Discovery | 5x10<sup>-8</sup> | 10 | 1 | 1 | 7** |
| Coffee consumption | Regular coffee cups consumed per day | Coffee and Caffeine Genetic Consortium 2015 [27] | 91,642 | Discovery | 5x10<sup>-8</sup> | 6 | 0 | 2 | 4*** |

**Table 5.**
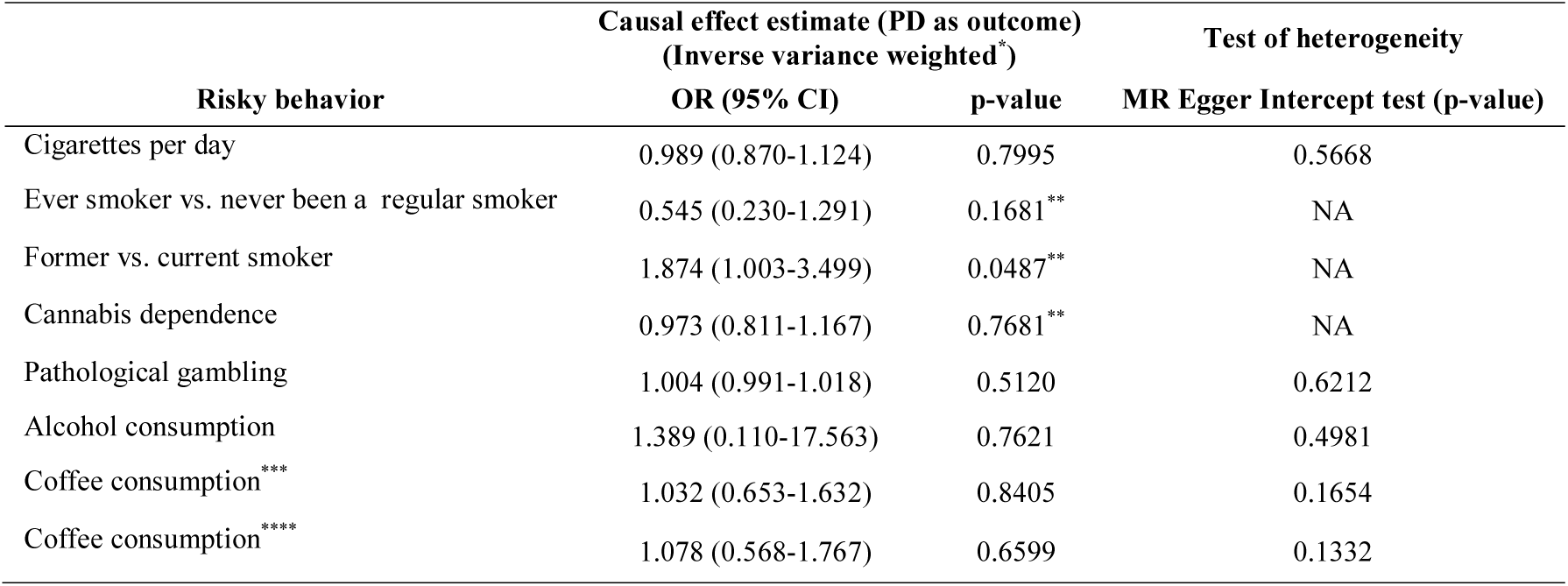
Causal effect estimates for habit-related behaviors from previously published GWAS. PD = Parkinson’s disease * Computed using second order weights **Computed using Wald estimate with delta method ***Excluding high LD SNP rs2470893 ***Excluding high LD SNP rs2472297

We further used a recently published GWAS on alcohol consumption using data from 112,117 individuals from the UK biobank ^24^. With seven genetic variants as a genetic instrument, we were able to replicate our finding of absence of causal association of alcohol consumption with PD (OR=1.389; 95%CI=0.110-17.563; p-value=0.7621). We further failed to observe any causal association of number of regular coffee cups per day with PD (OR=1.032; 95% CI=0.653-1.632; p-value=0.8405) ^27^.

We additionally investigated a protective causal role of cannabis dependence in PD by exploiting a recent GWAS study on cannabis dependence in 2080 cannabis-dependent cases and 6435 cannabis-exposed controls of European descent ^25^. The study reported a potential role of a cluster of highly linked 26 SNPs spanning a region on chromosome 10. The study further identified a putative functional SNP rs1409568 among this cluster responsible for the observed phenotypic association. Our investigation of a causal role of rs1409568 did not show evidence for a role of cannabis dependence in PD predisposition (OR=0.973; 95%CI=0.811-1.167; p-value=0.7681). A recent GWAS in 1531 Caucasians further reported absence of any significant SNPs with pathological gambling ^26^. We generated a genetic instrument based on the top hits from the study with a P-value cut off of <10^−4^ employing 45 uncorrelated genetic variants. We observed no causal association of PD with pathological gambling (OR=1.004; 95% CI=0.991-1.018; p-value=0.5120).

## Discussion

To the best of our knowledge, this is the first comprehensive study exploring the role of risky behaviors as causal factors for PD using a MR approach. The study suggests that the tendency-to-smoke trait is causally related to PD with individuals who started smoking being protected against PD. Our sensitivity analysis further demonstrated robustness of the reported association in the absence of any detectable pleiotropic effect. Furthermore, our secondary MR analysis did not show any causal association of other habitual traits including other smoking phenotypes such as number of cigarettes per day, cannabis dependence, pathological gambling, alcohol and coffee consumption with PD.

Numerous observational studies have previously shown an inverse association of smoking with PD. A meta-analysis merging smoking status trait from 33 different populations demonstrated a risk reduction by 36% for ever-vs never-smokers with consistent results in both case-control and cohort studies ^2^. Other epidemiological studies suggested significant gene-by-smoking interaction effects in PD^46, 47^. In our study, we observed a PD risk reduction of 31% for ever smokers vs never smokers. Although risk reduction effects demonstrated in an observational and in an MR study may not be comparable, a consistency in the direction of protective associations by both the approaches is an important finding.

To validate our results, we performed secondary MR analyses using other habit-related behaviors from other GWAS. The lack of association with a previously reported genetic instrument for ever smoker instrument as well as fomer smoker vs. current smoker may be explained by lower power of the GWAS with only one significant variant contributing to the instrument for both MR analyses. We also did not observe an association of PD risk and the number of cigarettes per day. One explanation would be that this continuous phenotype mainly just reflects the tobacco and nicotine exposure, whereas the ever vs never smoking might rather be a sign of risk taking behaviour. Our results thereby clearly imply the need for careful dissection of different smoking phenotypes. This will help understanding the causal role of the tendency to smoke on PD and reveal further insight into the development of the disease.

As outlined in the results section, our MR results on coffee consumption (cups per week) and alcohol consumption (drinks per week) also did not show significant causal associations. However, we cannot exclude that the analysis of coffee and alcohol consumption as quantitative traits may have the same limitations as the analysis of cigarettes smoked per day. Lastly, lack of a causal role of cannabis dependence observed in the present study needs to be further evaluated with stronger genetic instruments.

Absence of association with gambling in our analysis, however, could be attributed to the winner’s curse as SNP-exposure estimates used for calculation of casual estimates may be overestimated due to limited power of the study on gambling phenotype. Another important finding of our comprehensive MR analyses was absence of any causal role of drinks per week with PD. Using data from the UK biobank, we were able to replicate our finding of absence of a causal association of alcohol consumption with risk of PD. However, our study suggests a potential causal association of the number of sexual partners and PD risk. To the best of our knowledge, no epidemiological population-based study has yet examined the role of promiscuity on PD risk. Therefore, our MR results need to be interpreted cautiously and independent lines of validation of this association are required to confirm these results.

An important limitation of our current study is that we could not directly assess associations of individual genetic variants with potential confounders of association between risk behavior and PD due to the lack of knowledge of potential confounders and unavailability of individual-level data. Nevertheless, our sensitivity analysis demonstrated that exclusion of loci being associated with PD-associated phenotypes from the MR analyses had no effect on the overall association. We could not further provide data on the degree of sample overlap among GWAS datasets on exposure and outcome in our two-sample MR design. A considerable ovelap could bias the results towards the estimates generated through observational studies. However, this potential limitation could not have any impact on our results as the IVW method using second order weights employed in the current study is known to address this bias. And lastly, before drawing conclusions on the role of risky behavior on PD, we must recognize a critical limitation of our study that we could not do a stratified MR analysis based on dopaminergic treatment in cases as dopaminergic agonists are known to modulate risky behavior in PD patients.

Despite these limitations, to our knowledge, our study represents the most comprehensive MR study to date on risky behavior phenotypes and PD. An extensive sensitivity analysis including use of genetic instruments specific to individual phenotypic traits, use of previous studies, literature search for potential pleiotropic variants and brain expression analysis collectively demonstrate a strong causal protective role of smoking tendency on PD. Furthermore, the role of automobile speeding propensity as a causal risk factor emphasizes the need for a stratified MR based on dopamine-agonist treatment. The present study also demonstrates that careful interpretation of pleiotropic signals and sensitivity analysis based on biological function could lead to fine filtering of GWAS signals. Such an approach may assist in differentiating between mediators and exposures, thereby helping us to construct the causal pathways leading to PD^48^.

## Acknowledgments

We acknowledge the investigators of the original study^4^ and the PDGene database team^21^ for sharing the PD GWAS data used for this study.

## Author’s contributions

SG and CML conceived the idea of the study. SG designed the study, performed the data extraction from the literature and the statistical analyses, wrote the first draft, and revised the final draft of the manuscript. IK supervised study design and statistical analyses. FDG contributed to statistical analysis plan and conducted independent blinded statistical analyses. IK, FDG and CML participated in improving the design of the study, helped to draft the manuscript, and have been involved in revising the manuscript. MK and CK have been involved in revising the manuscript. All authors read and approved of the final manuscript.

## Availability of data and materials

Not applicable

## Ethics Approval and consent to participate

Not applicable

## Consent for publication

Not applicable

## Supplementary Table.

List of summary estimates used for the calculation of causal estimates for the primary MR analysis.

Exp: Exposure or phenotype

GRT: General risk tolerance

ADV: Adventurousness

ASP: Automobile speeding propensity

DPW: Drinks per week

ES: Smoking tendency (Ever vs never smoking)

NSP: Number of sexual partners

Chr: Chromosome

Pos: Position (GRCh37.p13)

EA (exp): Effect allele of the SNP in the exposure dataset

OA (exp): Other allele of the SNP in the exposure dataset

EAF: Effect allele frequency

Gene: Gene or nearby gene

β (exp): Effect estimate of SNP from regression analysis of the SNP with respective exposure

se (exp): Standard error of SNP from regression analysis of the SNP with respective exposure

p-value (exp): p-value of SNP from regression analysis of the SNP with respective exposure

EA (out): Effect allele of the SNP in the PD dataset

OA (out): Other allele of the SNP in the PD dataset

β (out): Effect estimate of SNP from regression analysis of the SNP with PD

se (out): Standard error of SNP from regression analysis of the SNP with PD

p-value (out):): p-value of SNP from regression analysis of the SNP with PD

PD = Parkinson’s disease

